# Hantaan virus polymerase has an *in vitro* Terminal Nucleotidyl Transferase activity that is conserved across the *Bunyaviricetes* class

**DOI:** 10.64898/2026.09.16.751791

**Authors:** Sergio Barata-Garcia, Quentin Durieux Trouilleton, Claire Debarnot, Hélène Malet, Juan Reguera

**Author notes:** To whom correspondence should be addressed. Juan Reguera.

## Abstract

**Abstract section:** *Bunyaviricetes* is a class of segmented negative strand RNA viruses (sNSV) that includes causative agents of severe zoonotic diseases resulting in hemorrhagic fever in humans and livestock. Hantaan virus (HTNV), family *Hantaviridae*, is an example of life-threatening bunyavirus transmitted by rodents which leads to Hantavirus hemorrhagic fever with renal syndrome in humans. Infection relies on the replication and transcription of their tripartite genomes through multifunctional RNA-dependent RNA polymerases (RdRp) in the context of ribonucleoproteins (RNPs) made of one viral RNA fragment, one RdRp and multiple copies of viral nucleoproteins. The RdRp, also known as L protein, performs replication by *de novo* initiation through a prime-and-realign mechanism while transcription initiates through a cap snatching mechanism, that engages a cap-binding domain and an endonuclease (EN) domain present in the L protein C and N terminal regions, respectively. In this work we show that HTNV L protein has a terminal nucleotidyl transferase (TNTase) activity over double-stranded RNA substrates (dsRNA) *in vitro*. We show that the TNTase activity is also present in other bunyaviral L proteins by comparing HTNV L protein activity with La Crosse (LACV, family *Orthobunyaviridae*) and Crimean Congo Hemorrhagic Fever Virus (CCHFV, family *Nairoviridae*) L proteins. Our results show that TNTase activity appears to be a general feature of Bunyaviruses and report the differences on substrate and nucleotide specificity for each family of virus.

**Importance:** The L proteins are preferential antiviral targets for the treatment of bunyaviral emerging diseases. Our results provide new ground for the characterization of L protein enzymatic activity *in vitro* which should be considered for the development activity assays, paramount for the screening of antiviral compounds. Indeed, TNTase activity can be easily misinterpreted as replication activity. We indicate here the key experimental setup and controls that would need to be considered to distinguish both activities. Our *in vitro* observations pave the way for future investivation of the potential TNTase role(s) in the infected cells. Proposed hypotheses concern potential functions in the preservation of genome integrity, the prevention of RIG-I innate immune response activation, or modification of RNA metabolism in the cell cytoplasm. The TNTase activity assays we describe here finally provide a straightforward biochemical approach for screening of inhibitors targeting RdRp activity.

## Introduction

Hantaan virus (HTNV) is a segmented negative-sense single-strand RNA virus (sNSV) that belongs to the *Bunyaviricetes* class, *Elliovirales* order, family *Hantaviridae* (1). HTNV is a rodent-borne virus, a widely distributed and prototype virus of Hantavirus, whose human infections cause hemorrhagic fever with renal syndrome reaching mortality rates up to 12% (2). It is thus a crucial target for the development of antivirals and prevention strategies to face future pandemic threats, as the recent outbreak of the related Andes hantavirus in a cruise ship in may 2026 (3–6). HTNV have a three-segmented genome consisting on small (S), medium (M), and large (L) segments, based on their size, which encode for the nucleoprotein (N), membrane glycoproteins Gn and Gc, and viral RNA-dependent RNA polymerase (RdRp) (also called L protein), respectively (7, 8).

The L proteins are multifunctional proteins responsible for both (i) viral RNA replication that initiates *de novo* in the absence of primer, and (ii) viral transcription that initiates through a cap-snatching mechanism characteristic of sNSVs (9, 10). Cap-snatching consists of holding a cellular capped RNA through the L protein cap-binding domain, prior to its cleavage by the L protein endonuclease (EN) domain and the subsequent use of obtained RNA capped oligonucleotide for the priming of the transcription reaction (8). Transcription and replication occur in the context of ribonucleoprotein (RNP) assemblies where each segment is coated by multiple copies of viral nucleoproteins and one L protein. *Bunyaviricetes* L proteins and influenza heterotrimeric polymerases specifically recognize their genomes by specific interactions with their 5′ and 3′ vRNA genomic ends which have specific binding sites in the protein surface. The vRNAs maintain a 5′-3′ distal duplex by complementary bases that is essential for replication and transcription initiation (11). The 5′ binding site recognizes the 5′ vRNA as a hook which is important for the regulation of the polymerase activity by allosterically activating the RdRp active site (10, 12). Two 3′ binding sites have been shown for sNSVs polymerases (13). The primary binding site is in the active site placing the 3′ template ready for *de novo* replication or cap-snatching transcription initiation. The secondary binding site for the vRNA 3′ end is at the surface of sNSV polymerases keeping the template’s ends attached to the polymerase during transcription and replication elongation. This RNA promoter binding strategy allows to keep the RNP circular throughout all the RNA synthesis process (14) (15) (16).

Enzymatic assays coupled to structural studies have been key for unveiling such sophisticated mechanism of genome processing by the RdRp of sNSV (10, 14, 17–19). Some species-specific behaviors highlight variations on this general functional trend. For instance, EN have been shown to be active or inactive *in vitro* and this is related to the presence or absence of a catalytic histidine in the EN active site (classified as His+ and His-ENs respectively) (20) (21). Is not clear how L proteins with His-ENs, such as those of arenaviruses and nairoviruses, perform cap-snatching. On another hand, Arenavirus L protein structure was reported together with biochemical assays showing a constitutively active polymerase in absence of 5′ vRNA (22) which contrasted with polymerase assays reporting the need of the dsRNA promoter with a non-templated 5′ G in the 5′vRNA (18), a non-templated nucleotide naturally present in arenavirus genomes. In arenaviruses the polymerase activity is regulated by the small viral protein Z, which is unique to this viral family (23). Thus, further studies on replication and transcription activities are required for the precise comprehension of L protein activities and their regulation for each particular bunyaviral family.

All viruses with an RNA genome face the challenge of maintaining the genome integrity which may be affected by cellular RNA modifications or high RdRp error rates. Most Bunyavirus and Orthomyxovirus perform a prime-and-realign mechanism during RNA synthesis initiation allowing for the addition of terminal repeats at the end of the genome if its correct length is compromised (9) (24) (Fig. 1). Arenaviruses even incorporate a non-templated G in the 5′ end of their genomes, a prime-and-realign associated phenomena. However, other viral RdRps follow different strategies for maintaining genome integrity. Particularly, TNTase activity has been reported as a common behavior of viral RdRps. For example, Wuhan nodavirus (WhNV) TNTase activity repairs the 3′ initiation site restoring replication initiation *in vitro*, as a protection mechanism of the 3′ template vRNA against cellular exonucleases (25). Among others viral RdRps with TNTase activity we find Sapovirus 3Dpol which has CTP-preference incorporation (26); Hepatitis C virus (HCV) with UTP preferential incorporation (27) or Nodovirus 3Dpol (28) with broader TNTase activity. Besides, TNTase studies of flavivirus HCV and bovine viral diarrhea virus (BVDV) RdRps revealed that changes in template sequences and/or structure could affect the efficiency and preference of nucleotide incorporation (29). To our knowledge, no TNTase enzymatic activity has been reported for *Bunyaviricetes* L proteins yet.

**Figure 1.**
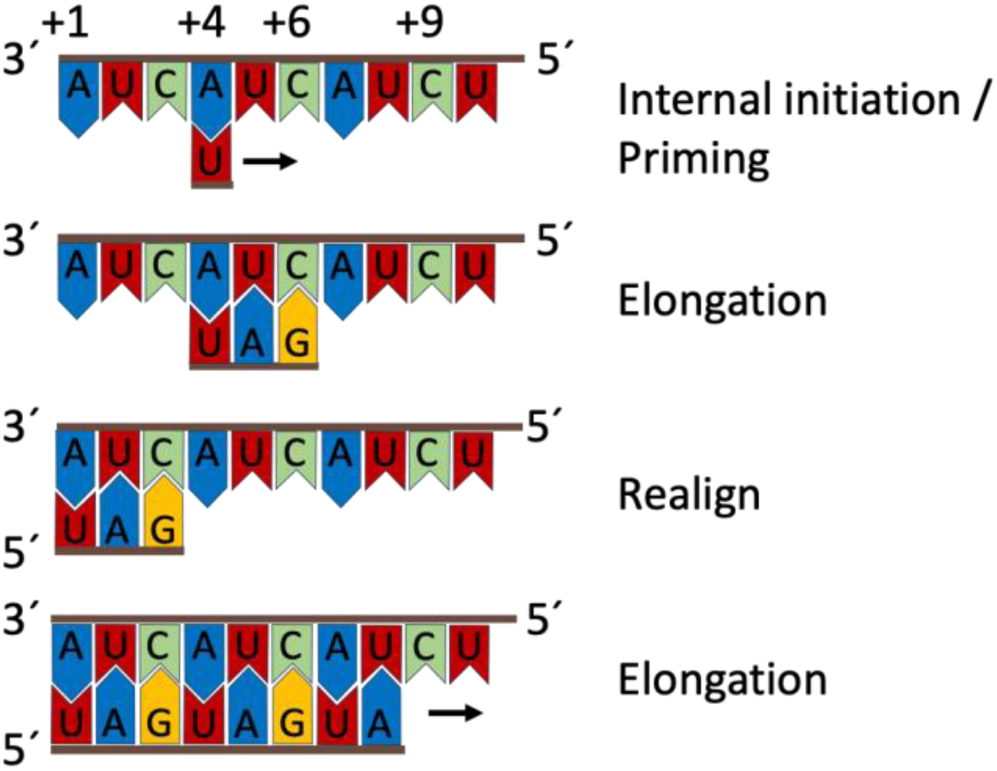
Scheme of the prime-and-realign RNA synthesis initiation characteristic of bunyaviruses and orthomyxoviruses. The HTNV genome sequence is used. The polymerase starts the RNA synthesis in the second triplet of the genome for *de novo* initiation of genome replication. After the synthesis of three nucleotides the RNA is realigned upstream for subsequently proceed with the RNA synthesis elongation.

In this work, we characterize the enzymatic activities of the L protein of HTNV virus as the EN and RdRp activities, and reveal the existence of a TNTase activity. We study the nucleotide and substrate specificity of incorporation. In addition, we extend this finding to other two L proteins from the *Bunyaviracetes* class: (i) La Crosse virus (LACV) L that belongs to the *Elliovirales* order and the *Peribunyaviridae* family and (ii) Crimean Congo Hemorrhagic Fever Virus (CCHFV) L that belongs to the *Hareavirales* order and the *Nairoviridae* family. This finding is relevant for understanding the *in vitro* activity of these enzymes and the proper interpretation of biochemical assays on bunyaviral L proteins. In addition, by EN assays performed in parallel with the three L proteins we confirm the HTNV and LACV (His+) L proteins as active and the CCHFV (His-) as inactive. Our results have implications for the understanding of L protein’s molecular mechanisms of replication and possible interactions with host RNAs.

## Methods

### Expression and purification of L proteins

Codon-optimized gene coding for HTNV L (GenBank reference: NC_005222.1, UniProtKB reference: P23456) and CCHFV L (GenBank ref: NC_005301.3, Uniprot ref: Q6TQR6) carrying N-terminal His-tag followed by TEV protease cleavage site (MGHHHHHH6xHis-tagDYDIPTTENLYFQTEVG) were cloned into pFastBac™1 donor plasmids. Codon-optimized gene coding for LACV strain LACV/mosquito/1978 L protein sequence (GenBank code: EF485038.1, UniProt code: A5HC98) was cloned without a tag in pFastBac™1. The HTNV L cap-snatching EN domain (D97A) and RdRp active site (HTNV L_SDD1097AAA_) mutants were generated by Quick-change site-directed mutagenesis. All recombinant baculoviruses were produced using DH10BACY *E. coli* competent cells and recombinant proteins were expressed in *Trichoplusia ni* High 5 cells (Invitrogen PN/51-4005 lot 1783124). For HTNV L, infected cells were harvested at 96 h post-infection by centrifugation (10 min, 4 °C, 5000xg). Following resuspension in buffer A (30 mM Tris-HCl pH 8.0, 300 mM NaCl, 5% glycerol, 2mM TCEP) supplemented with 10 mM imidazole, protease inhibitor cocktail tablets (Roche), MgSO_4_, DNAase I and RNAse A, cells were disrupted by sonication on ice and centrifuged at 21.000xg for 45 min at 6°C. The clarified supernatant was directly loaded into a Ni-NTA affinity column (HisTrap™ FF crude, GE Healthcare) and washed with buffer A supplemented with 40 mM imidazole. HTNV L was subsequently eluted with buffer A supplemented with 250 mM imidazole. HTNV L fractions were pooled and imidazole concentration was diluted by 2 using buffer A before loading on heparin column (GE, HiTrap™ Heparin HP). Washing step in 9% buffer B containing 30 mM TRIS-HCl pH 8.0, 300 mM NaCl, 5% glycerol, 2 mM TCEP, 1M NaCl was followed by an elution with 30 % of buffer B. HTNV L fractions were concentrated (30-kDa-cutoff Amicon Ultra unit) and further purified by size exclusion chromatography (GE, Superose 6 increase 10/300 GL) with buffer C (30 mM Tris-HCl pH 8.0, 500 mM NaCl, 5% glycerol, 5 mM TCEP). Peak fractions analyzed by SDS-PAGE 6% were concentrated and flash-frozen in liquid nitrogen and stored at -80 °C for further biochemistry studies.

Untagged LACV L protein was produced as described for HTNV-L protein. High 5 infected-cell pellets were resuspended in buffer A (50 mM Tris-HCl pH 8.0, 500 mM NaCl, 5 mM beta-mercaptoethanol (BME) and purified by sequential ammonium sulfate precipitation, isocratic gradient elution in heparin column with buffer B (50 mM Tris-HCl pH 8.0, 250 mM NaCl, 5 mM BME) supplemented with 1M NaCl and SEC (Superose 6 increase 10/300). Pooled LACV L in 50 mM Tris-HCl pH 8.0, 150 mM NaCl, and 5 mM BME was flash-frozen in liquid nitrogen and stored at -80 °C for further biochemistry studies. CCHFV L polymerase was purified following the same conditions as the HTNV L recombinant proteins.

Pure proteins analyzed by SDS/PAGE 6% were processed by mass spectrometry facility of Marseille Proteomics (MaP-CRCM). Briefly, gel bands were cut, destained in 100 mM NH_4_HCO_3_ in 50% acetonitrile, dried at room temperature, then rehydrated. Cysteines were then reduced using 10 mM DTT in 100 mM NH_4_HCO_3_ pH 8.0 for 45 min at 56°C before alkylation in the presence of 55 mM iodoacetamide in 100 mM ammonium bicarbonate pH 8.0 for 30 min at room temperature in the dark, and bands were finally washed twice with 25 mM NH_4_HCO_3_ pH 8.0 and digested with high-sequencing-grade trypsin (Promega, Madison, WI). Mass spectrometry analysis was carried out by LC-MSMS using Orbitrap Mass Spectrometers (Thermo Electron, Bremen) online with a nanoLC Ultimate 3000 chromatography system (Thermo Fisher Scientific™, San Jose, CA).

### Endonuclease assay

Endonuclease activity in the full-length L protein context was measured by degradation of fluorescent-RNA substrate. Nuclease activity assays were carried by mixing 2 μM of full-length L protein (HTNV L, LACV L or CCHFV L) or 5 μM Toscana virus (TOSV, *Phenuiviridae* family) EN domain (as described in Jones et al. 2019 (21)) with 0.5 μM GA-rich RNA (5′-FAM labeled) in a buffer containing 30 mM Tris-HCl pH 8.0, 500mM NaCl and 5% Glycerol and 4 mM TCEP for HTNV L or 5 mM BME for CCHFV L and LACV L. Reactions were supplemented with 2 mM Mn^2+^, Mg^2+^, EDTA and 0.5 mM DPBA where indicated. TOSV EN was used as a positive control in all experiments with a buffer containing 35 mM Tris-HCl pH 7.5, 150 mM NaCl and 2 mM TCEP in presence of 2 mM Mn^2+^ and 0.5 mM DPBA where indicated. After an initial screening performed at 30 °C for 30 min, 1, 3 and 6 h, further experiments were performed at 30 °C for 1h. The reactions were stopped by loading buffer addition (Tris–borate–EDTA, 8 M urea, 5% glycerol). Samples were pre-heated at 70 °C for 5 min and placed on ice for 5 min before loading onto 20% Poly-acrylamide (PPA) 8 M Urea gels. RNA migration on PPA-Urea gel was performed in 0.5x TBE buffer (89 mM Tris-borate, 2 mM EDTA at pH 8.3) at 250 V for 1h. Gels were visualized with a Phosphorimager Typhoon system and analyzed with ImageQuant TL program (Amersham).

### Terminal transferase activity assay

For TNTase activity, RNA substrates at 5 µM were preheated at 65 °C for 5 min and cooled down on ice for 5 min. 0.5 µM of L protein was then mixed with RNA substrates in a reaction buffer containing 30 mM Tris-HCl pH 7.5, 150 mM NaCl, 30 mM KCl, 2 mM TCEP, 0.1 mg/ml BSA for HTNV L, 50 mM Tris-HCl pH 7.5, 150 mM NaCl, 30 mM KCl, 5 mM BME, 0.5 mg/ml BSA, for LACV L, and 50 mM Tris-HCl pH 7.5, 150 mM NaCl, 30 mM KCl, 2 mM TCEP, 0.1 mg/ml BSA for CCHFV L. The reaction was also supplemented with 5 mM Mg^2+^ and 0.6 U µl^-1^ RNAase inhibitor. TNTase reaction was started by adding to the mix 0.2 µCi/µl of the indicated [^32^P-NTP] and 0.04 mM of the same cold NTP. When *in vitro* RNA synthesis was studied, reactions were started by adding 0.5 mM GTP/CTP/ATP, 0.04 mM UTP, and 0.2 µCi/µl [^32^P-UTP]. The resulting mixes were incubated at 30 °C for 3 h. All reactions were stopped by adding 2x loading buffer, heating 10 min at 70 °C and were resolved on a 20% TBE polyacrylamide gels-7 M Urea. The RNA products were separated with 1x TBE buffer for 2h30min. The gels were exposed on a storage phosphor screen and developed with an Amersham Typhoon scanner.

All synthetic RNA oligos used for activity experiments were chemically synthesized by *Microsynth AG*. RNA oligos smaller than 9 nucleotides were acquired from GE Healthcare Dharmacon. *In vitro* RNA synthesis was studied by radiolabeled [α-^32^P] NTP incorporation in RNA product. Unlabeled synthetic RNA oligos shown in Table 1 corresponding to viral RNAs (vRNA) were used as substrates. As molecular size marker, we radiolabeled specific RNA with [γ-^32^P] ATP by mixing 0.5 µl T4 Polynucleotide Kinase (T4 PNK) with 1x PNK buffer (NEB), 10µM vRNA and 0.5 µCi/µl [γ-^32^P] ATP in a final volume of 10 µl at 37 °C for 10 min, followed by 10 min at 70 °C to stop the reaction and addition of loading buffer.

**Table 1.**
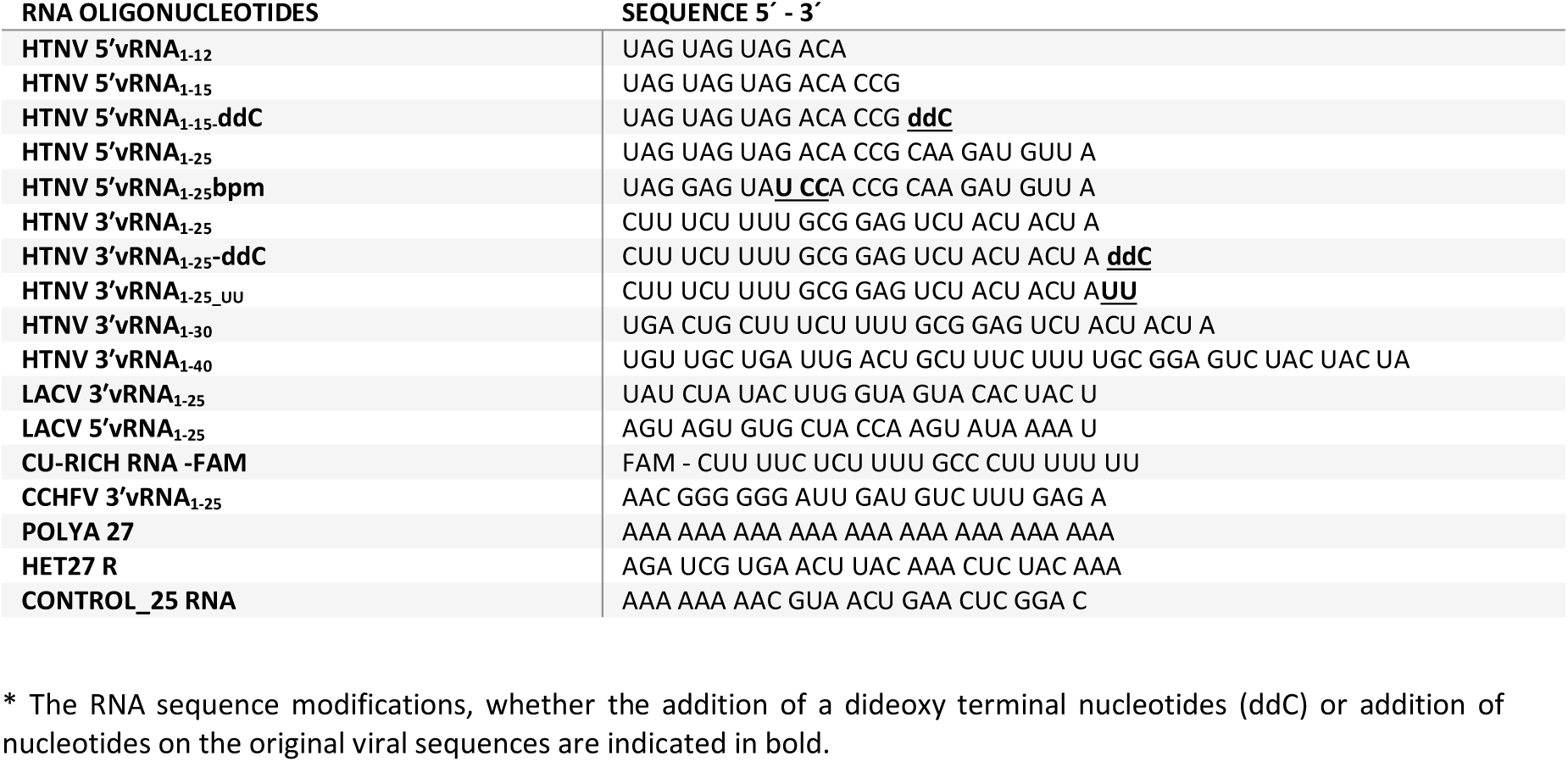
oligonucleotides RNA.

HTNV, LACV and CCHFV 5′ and 3′ genomic end RNAs were predicted using Mfold within UNAfold package web server. Computer-predicted RNA fold was carried out using the default parameters and output RNA secondary structures were consider to create the RNA fold figures (Supplementary fig. 2).

## Results

### Recombinant production of bunyaviral L proteins

We used the baculovirus expression system in insect cells and codon-optimized genes for the overexpression of the N terminal 6xHis tagged HTNV L and CCHFV L proteins and the untagged LACV L, since the tag in LACV L protein has been shown to strongly diminish polymerase activity (14). HTNV and CCHFV L proteins were purified following three steps of chromatography as described in methods (Fig. 2A, 2C). For the untagged LACV L protein we substituted the affinity chromatography step with an ammonium sulfate precipitation (Fig. 2B). In all cases, we obtained homogeneous protein of the expected size (246 kDa for HTNV L, 263 kDa for LACV L and 447 kDa for CCHFV L) and the protein identity was confirmed by mass spectrometry (see methods). The protein yields after expression and purification was of 0.5 mg/L of culture for HTNV L, 0.8 mg/L for LACV L and 0.3 mg/L for CCHFV L. Mutants of the cap-snatching EN active site (HTNV L_D97A_) and of the RdRp active site (HTNV L_SDD1097AAA_) could be generated and L proteins could be purified with yields similar to the ones obtained for HTNV L wild type (HTNV L_WT_). In conclusion, the expression and purification of L proteins in baculovirus systems is possible for a wide number of bunyaviral L proteins following similar purification procedures.

**Figure 2.**
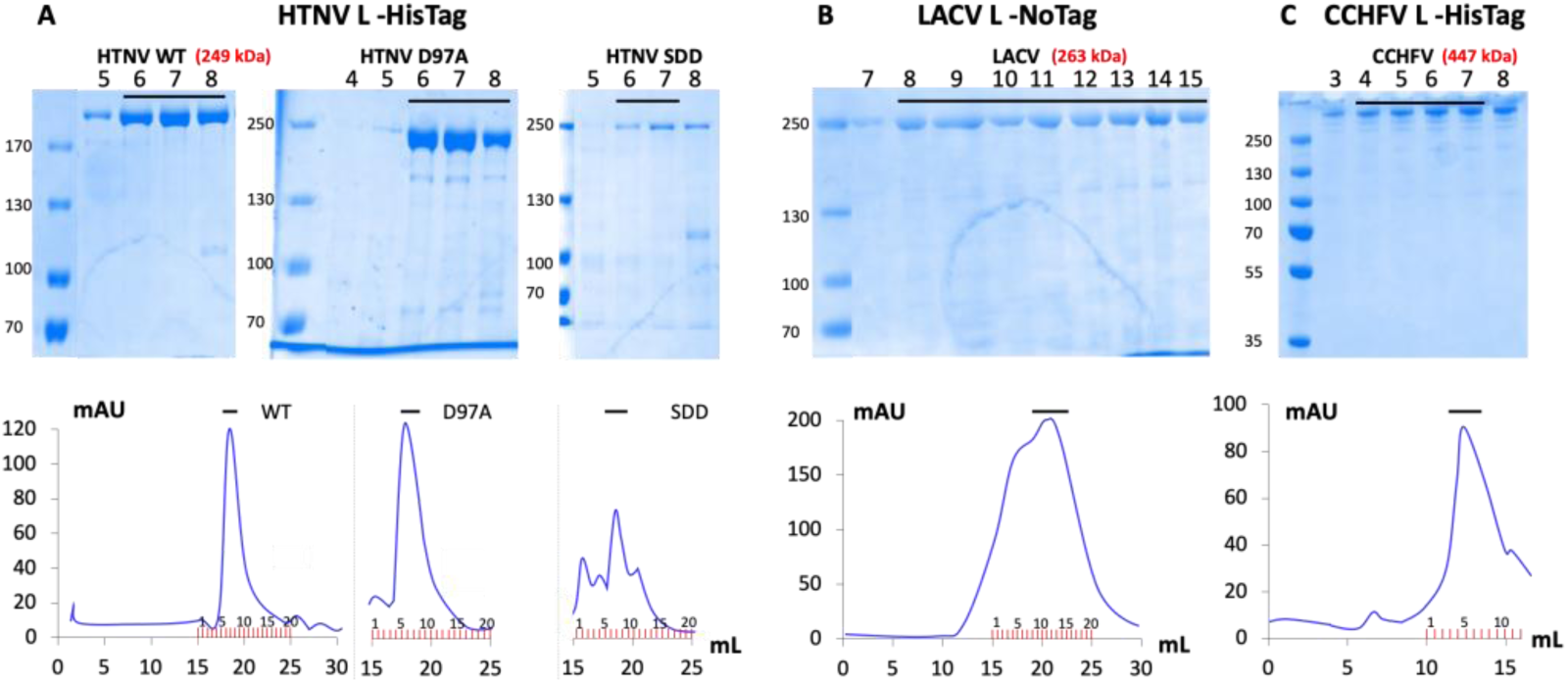
Purification of recombinant Full-Length L proteins. SDS-PAGE gel of L protein fractions after size exclusion chromatography (Superose 6 increase 10/300) for HTNV L – HisTag (A), LACV L – NoTag (B) and CCHFV L – HisTag (C). The size exclusion chromatography profile of each gel is displayed below. The molecular weight ladder is shown on the left of each gel. The pooled fractions for the final protein preparation are indicated with a black line. L protein molecular weights are indicated in red. Molecular weight markers and lanes belong to the same gel.

### Full-length L protein endonuclease activity assays

The cap-snatching ENs of L proteins exhibit metal ion-dependent activity and have been extensively characterized, particularly within isolated domains of Bunyavirus and Influenza Orthomyxovirus. (20, 21, 30–33). The EN activity of the full-length L proteins has been less characterized *in vitro* for viruses of the *Bunyaviricetes* class. We thus explored the cation-dependent EN activity of the full-length L proteins of HTNV, LACV and CCHFV, including the Toscana virus (TOSV, family *Phenuiviridae*) EN domain as positive control.

Endonuclease activity assays were performed with the four proteins (Fig. 3A). In the absence of metal ions, TOSV and LACV ENs showed some basal activity. This has been seen before for active bunyaviral ENs indicating traces of metal ions or perhaps traces of contaminant RNAses in the buffer. Indeed, LACV L, purified without tag affinity chromatography, showed some basal activity in presence of EDTA chelator, indicating the presence of an RNAse contamination background. In presence of Mg^2+^ and Mn^2+^ metal ions, LACV and HTNV L proteins showed a strong activity which could be inhibited by the 2,4-dioxo-4-phenylbutanoic acid (DPBA) inhibitor, a well known inhibitor for cap-snatching ENs which chelates both metal ions in the active site (21) (30) (20). In contrast, CCHFV L showed no activity at all in the same experimental conditions. As expected, the TOSV EN control was active in presence of Mn^2+^ but not Mg^2+^. Similar behavior was found for HTNV and LACV ENs previously (20). The Mg^2+^ dependent activity shown here by full-length LACV and HTNV L proteins evidences that Mg^2+^ can be used for cap-snatching *in vivo*, even if only Mn^2+^ is effective on isolated ENs. This is consistent with what was reported for influenza heterotrimeric RdRp (34). However, the absence of activity for CCHFV L with Mg^2+^ and Mn^2+^ metal ion poses the question of how this protein carries out the cap snatching transcription initiation. Arenavirus full-length L proteins, with a His – EN, have been reported to have a behavior similar to CCHFV L protein (18).

**Figure 3.**
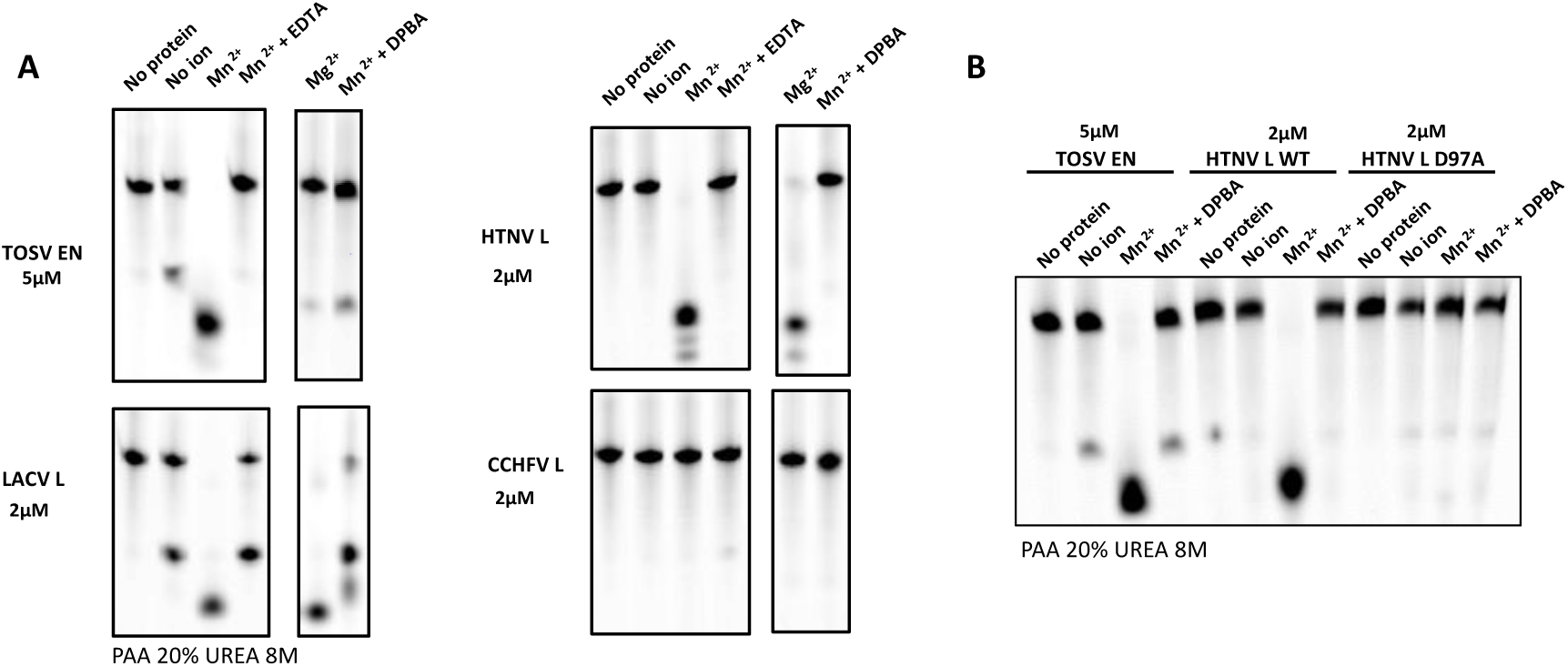
Endonuclease activity assays. **(A)** Comparative assay of EN activity in the full-length context for HTNV L_WT_, LACV L_WT_ and CCHFV L_WT_ versus TOSV EN domain. The activity is shown in presence or absence of Mn^2+^, Mg^2+^, (2,4-dioxo-4-phenylbutanoic acid) DPBA inhibitor and EDTA. **(B)** EN activity of full length HTNV L_WT_ and HTNV L_D97A_. TOSV EN domain is used as a positive control. Reactions are performed with the protein concentration indicated at 30 °C for 1 h with a fluorescence-labeled GU-rich RNA. Reaction products were resolved on 20% PAA 8 M urea gels.

To further associate the detected endonuclease activity to the EN domain of HTNV L, we carried out site-directed mutagenesis of Asp97 (D97), involved in the two-metal ion coordination, to alanine (D97A) and analyzed the effect on catalytic activity (Fig. 3B). We see, in the same experimental conditions, the same behavior of TOSV EN and HTNV L_WT_ but a total lack of activity of HTNV L_D97A_ which provides further evidence that the endonuclease activity detected for HTNV is directly caused by the EN domain of HTNV L. This results are consistent with previously reported activity assays (35, 36).

### HTNV L protein exhibits a Terminal Transferase activity (TNTase) *in vitro*

In order to assess the RdRp activity of the HTNV L protein we carried out replication assays by incubating the L protein with its 25 nucleotides long 3′ vRNA template (3′vRNA_1-25_) and the 15 nucleotides long 5′ vRNA (5′vRNA_1-15_) allosteric regulator using MgCl_2_ and NTP substrates and [α-^32^P]-UTP to detect the newly synthesized RNAs (Fig.4). As a negative control of replication, we performed the reaction in presence of UTP only, in the absence of the other NTPs. We unexpectedly found in this condition a very strong incorporation of [α-^32^P]-UTPs into the 3′vRNA_1-25_ and the 5′vRNA_1-15_, respectively (Fig. 4A, lane 1), as evidenced when comparing with the size of the 3′vRNA_1-25_ and 5′vRNA_1-15_ labeled with [γ-^32^P]-GTP (Fig. 4A, marker panel). One to four [α-^32^P]-UTPs were added, with a preference of a two-nucleotide addition. When we substituted the 15 nucleotides long 5′vRNA by a 12 nucleotide long vRNA, we could see the same product of the 3′vRNA length and a shorter product consistent with the incorporation of two [α-^32^P]-UTP in the 5′ 12 nucleotides long RNA (Fig. 4A, lane 2). The visualized addition of terminal nucleotides on both the 3′ and 5′vRNA corresponds to a terminal transferase activity (TNTase). Importantly, it can easily be misinterpreted as replication activity. Indeed the same pattern of RNA products can be seen when incubating with UTP only (see Fig. 4A, lane 1 – conditions for TNTase activity) or all the NTPs (Fig. 4A, lane 3 – conditions for replication activity), with slightly lower intensity in presence of all the NTPs. Increasing the size of the 3′ RNA to 30 and 40 nucleotides long also detected RNA that would be consistent in size to either TNTase or *de novo* RNA synthesis. However, in none of the reactions with all NTPs present we could find additional intermediate products characteristic of *de novo* RNA synthesis. The absence of replication signal in presence of 3′vRNA_1-25_ and 5′vRNA_1-15_ can be explained by the complementarity of the 3′ and 5′vRNA that form double-stranded RNA and prevents binding *in vitro* as single stranded RNA, like has been previously shown in (10). These results are consistent with the evidence that *in vitro* replication with small vRNA can only be observed when the 5′-3′ dsRNA duplex is partially disrupted at the 5′ and 3′ extremities for instance by mutagenesis (14) (10) (Supplementary Fig. 1). This is necessary *in vitro* to allow the 5′ vRNA nt 1 to 11 to bind as a single-stranded hook in the 5′vRNA binding site and activitate the RdRp, while the 3′vRNA extremity can, as single-stranded RNA, pass through the template entry tunnel to reach the polymerase active site (10).

**Figure 4.**
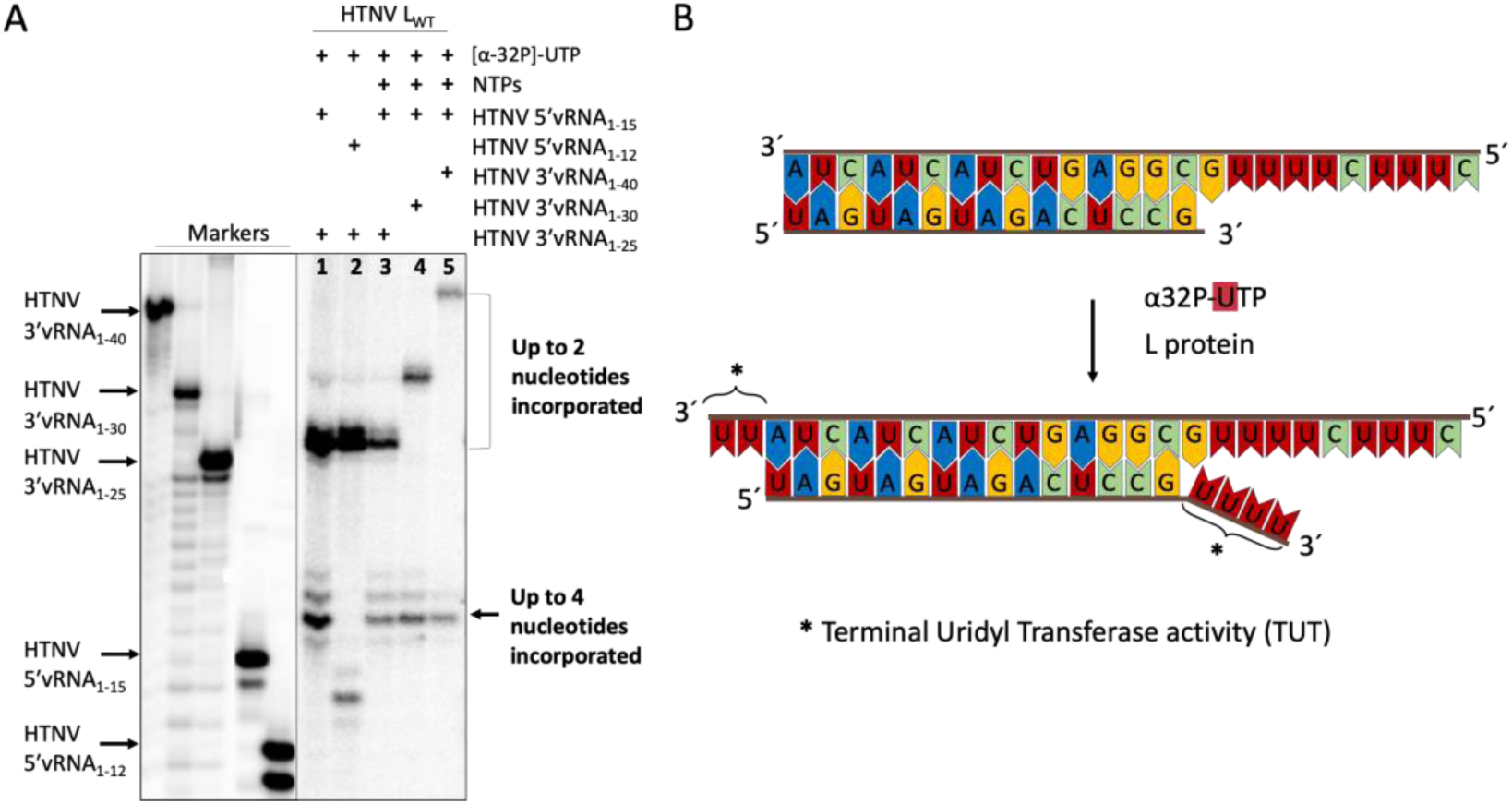
Terminal transferase activity of HTNV L_wt_. **(A)** Size markers are shown on the left panel and correspond to vRNAs labeled with [γ-^32^P]-GTP. On the right panel, TNTase activity of HTNV L_WT._ Lanes 1 and 2 are obtained by incubating HTNV L_WT_ with UTP only. Lanes 3 to 5 are obtained by incubating HTNV L_WT_ with the four NTPs. In all cases UTP is radiolabeled. Molecular weight markers and lanes belong to the same gel. **(B)** Schematic interpretation of the TUTase . Double stranded 3′/5′vRNA in presence of HTNV L_WT_ and UTP generates the extension of 2 and 4 nucleotides in the 3′OH RNA of both strands (*).

Altogether these results indicated that we do not visualize replication activity when HTNV L is incubated with both its wild type 5′ and 3′ vRNA ends, but that we instead identify a strong TNTase activity (Fig. 4B).

### HTNV L protein specifically incorporates UTP in the 3′ vRNA blunt end of its doubled-stranded vRNAs

Considering the high complementarity of the 3′ and 5′ vRNAs, we wondered if the TNTase activity is depending on (i) the 3′ vRNA sequence, (ii) the 5′ vRNA sequence or (iii) the formation of a double-stranded hybridized form of the 3′ and 5′vRNA. To test this, we incubated HTNV L with either (i) the 3′vRNA_1-25_ alone, (Fig. 5, lane 1), (ii) the 5′vRNA modified at the 3′ with a dideoxyCMP (5′vRNA_1-15ddC,_ Fig. 5, lane 2), or (iii) both the 3′vRNA_1-25_ and 5′vRNA_1-15ddC_ (5′vRNA_1-15ddC,_ Fig. 5, lane 3 to 6, with increasing concentration of 5′/3′ RNA duplex in presence of radioactive UTP). 5′vRNA_1-15_ddC was chosen as the absence of a 3′OH at its 3′end prevents the elongation of this RNA on the 3′vRNA template by incorporation of non templated nucleotides. No significant signal is visualized when HTNV L is incubated with either the 3′vRNA_1-25_ or the 5′vRNA_1-15ddC_ (see Fig. 5, lanes 1 and 2), indicating that the presence of double-stranded form of the 3′ and 5′vRNA is necessary. In presence of both 3′vRNA_1-25_ and 5′vRNA_1-15ddC_ the TUTase activity was visualized only on the 3′vRNA_1-25_ (Fig. 5, lanes 3 to 6), whereas it was observed on both 3′vRNA_1-25_ and 5′vRNA_1-15_ in the absence of the ddC modification of the 5′vRNA_1-15_ as shown above (see Fig. 4A lane 1). Altogether this showed that the HTNV L was able *in vitro* to incorporate UTP in the 3′ vRNA blunt end of vRNAs when they are in their double-stranded hybridized form (see Fig. 4B).

**Fig. 5.**
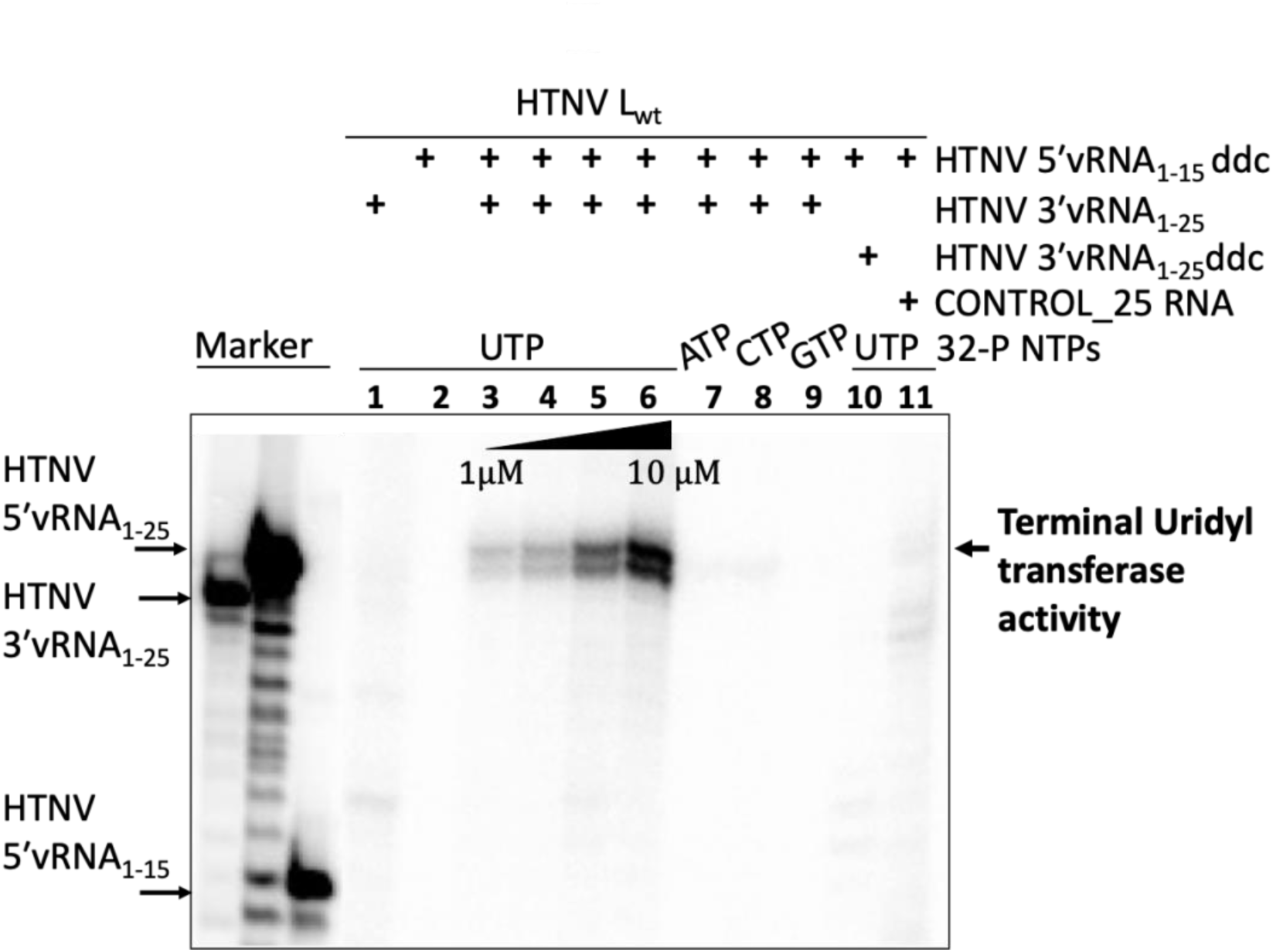
Terminal transferase activity analysis of HTNV L_wt_ on the 3′vRNA 1-25 template. **(A)** The single NTP used in each reaction is indicated. The concentration of RNA used in the reaction is 10 μM, except in lines 3 to 6 where it is 1, 2, 5, and 10 μM, respectively. No significant TNTase activity is detected when incubating HTNV L_wt_ with 3′vRNA_1-25_ or 5′ vRNA_1-15_ alone (lanes 1-2). Increasing concentrations of 5′/3′ vRNA show specific TUTase activity at the 3′vRNA (lanes 3 – 6). Non-significant incorporation of ATP, CTP, and GTP is detected (lanes 7 – 9). No TNTase activity is detected in the presence of dideoxy-C in the 3′vRNA (lane 10). Light background activity is shown on an control-25nt long RNA incubated alone with HTNV L_wt_ (lane 11). Size markers are indicated on the left by RNAs labeled with [γ-^32^P] ATP.

In order to determine the specificity of nucleotide incorporation on viral RNA we carried out a set of reactions with radiolabelled [α-^32^P]-ATP, UTP, GTP and CTP, incubating HTNV L_WT_ with 3′vRNA_1-25_ and 5′vRNA_1-15ddC_. When we performed the reaction with the other radiolabelled NTPs alone, we did not observe a similar incorporation of any of them with respect to UTP (Fig. 5, lanes 7-9). We included a 3′vRNA_1-25_ddC as a negative control to confirm that, as expected, nucleotides are added to the 3′ ends of the RNA (lane 10). Only a light background signal was detected with a control-25 RNA with no sequence similarity to the Hantaan promoters (lane 11) (see Table 1). Altogether these results showed that HTNV L has a specific Terminal Uridyl Transferase (TUTase) activity that incorporates two uridines at the 3′ of the 3′-OH end of its dsRNA promoter.

### HTNV L protein TUTase activity is specific of double-stranded RNA

We next wanted to determine the substrate specificity of HTNV L_wt_ TUTase activity. We first tested if additional UTP could be incorporated during subsequent incubations by using a modified 3′vRNA_1-25_ with two extra UMP at 3′end (3′vRNA_1-25_UU) and the 5′vRNA 1-15 (Fig. 6, lane 7). This further extended 3′vRNA_1-25_UU, generating an RNA that is elongated by 5 Uridines compared with unmodified 3′vRNA_1-25_ substrate (Fig. 6, lanes 5-6). In contrast, the incubation of HTNV L_wt_ with 3′vRNA_1-25_, 5′vRNA_1-15_ddC or 3′vRNA_1-25_UU alone resulted in no product formation, as expected (Fig. 6, lanes 1-4). This leads to the conclusion that double-stranded RNA with two U incorporated can be further extended with polyU by subsequent incubations with HTNV L_WT_. Mutation of the RdRp catalytic site (HTNV L_SDD1097AAA_) abrogated the TUTase activity on the HTNV dsRNA substrate confirming that the observed activity occurs in the HTNV L RdRp active site (Fig. 6, lane 8).

**Figure 6.**
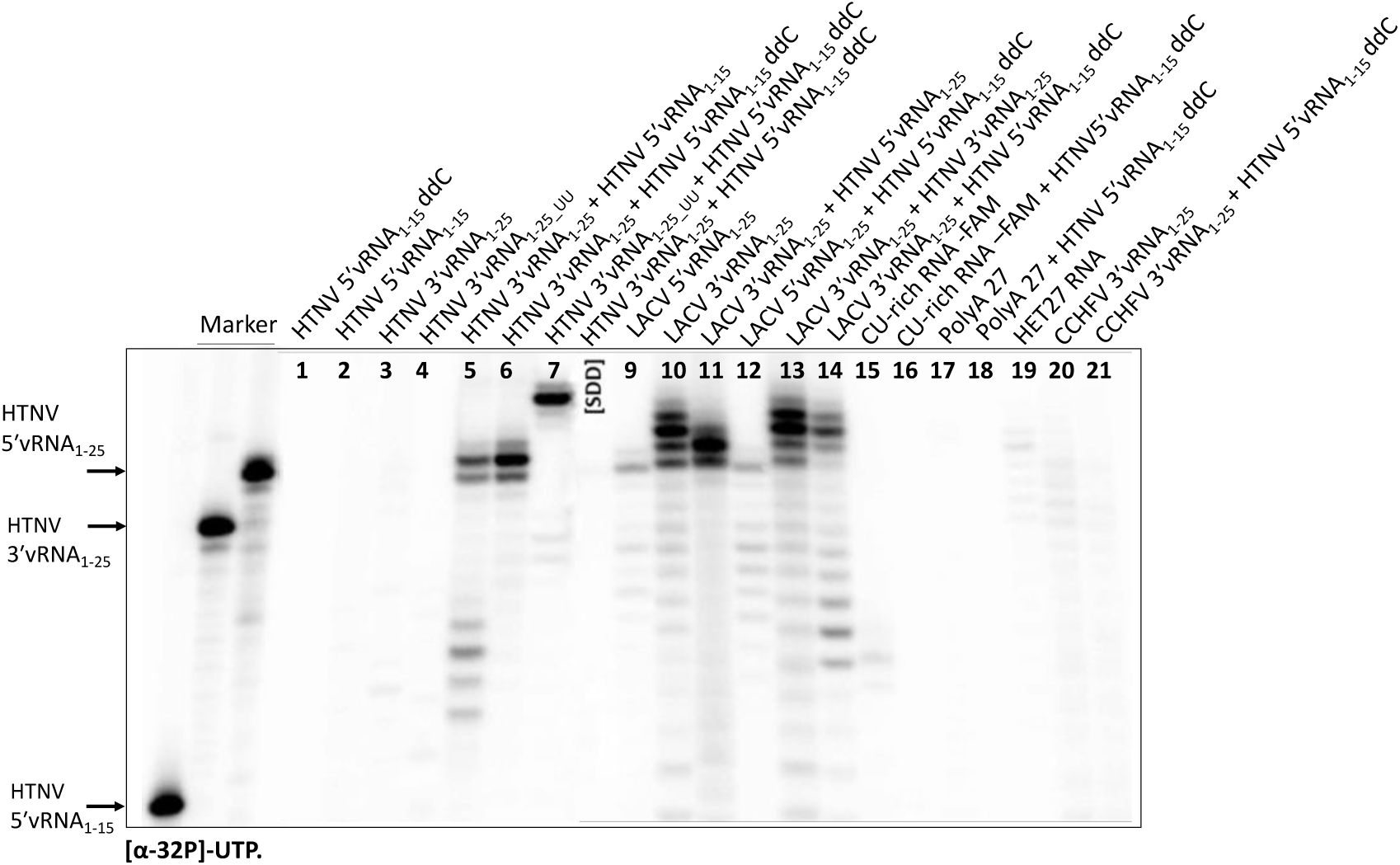
TUTase activity of HTNV L_wt_ with different substrates. TUTase reactions were carried out only with [α-^32^P]-UTP. Size markers are indicated on the left by vRNAs labeled with [γ-^32^P]-ATP. No activity was observed with HTNV and CU-rich or polyA single stranded RNAs (ssRNAs) (lanes 1 – 4 and 15 – 18). UMP incorporation is observed with ds vRNA of HTNV and both ss and ds vRNA of LACV (lanes 3 – 7 and 9-11). The addition of HTNV 5′vRNA_1-15_ ddC does not increase the TUTase activity on LACV control substrate (lanes 14, 16, 18, 21). The active site mutant (HTNV L_SDD1097AAA_, indicated as SDD) totally abolishes the activity (lane 8).

We also wondered if the TUTase activity was specific to HTNV vRNA or if this activity could also be visualized when incubating HTNV L_WT_ with other bunyaviral vRNA or with random RNAs. Surprisingly, we observed a clear activity over LACV 3′vRNA_1-25_ in comparison with the almost undetectable activity of LACV 5′vRNA_1-25_ RNA incubated alone (Fig. 6, lanes 9 – 10). RNA secondary structure prediction (see methods) showed that LACV 3′vRNA_1-25_ tends to form a blunt end with an extra U in the 3′ through a pan-handle structure with an 8 nucleotides dsRNA region (Supplementary Fig. 2). This is not the case for the LACV 5′vRNA_1-25_ or the other RNAs tested (see Fig. 6, lane 9 and 15-21) which, are not uridylated by HTNV L_WT_. The LACV 5′/3′ dsRNA (Fig. 6, lane 11) yielded similar products as the HTNV 5′/3′ dsRNA (see Fig. 6, lane 5) with even an increased activity. Considering the activation of HTNV L_WT_ by 5′ vRNA to initiate replication activity (10), we wondered if this allosteric regulation would have an effect on TUTase activity. Thus, we incubated the protein with HTNV 5′vRNA_1-15_ddC and LACV 3′vRNA_1-25_ or LACV 5′vRNA_1-25_ (Fig. 6, lane 12 and 14). We do not observe strong changes in the activity compared to the same conditions without the HTNV 5′vRNA_1-15_ddC (see Fig. 6, lane 9 and 11) but perhaps an enhanced RNA degradation. Likewise, taking into account the strong activity with LACV 3′vRNA_1-25_ alone, we added HTNV 3′vRNA_1-25_ to the reaction in order to study whether it is able to compete through specific interactions with HTNV L_WT_ (Fig. 6, lane 13), however no significant differences were observed.

Altogether, these results lead us to the conclusion that HTNV L_WT_ presents a 3′ TUTase activity by the RdRp active site which is specific for blunt-ended or few nucleotides overhang dsRNAs and is not significantly affected by the allosteric regulation of the 5′ vRNA.

### Conservation of bunyaviral L protein terminal nucleotidyl transferase activity

Intrigued by the TNTase activity detected for the HTNV L, we wondered if this activity was a general trend for bunyaviruses. To investigate this, we tested the potential TNTase activity of two additional L proteins from LACV and CCHFV, representatives of the *Peribunyaviridae* and *Nairoviridae* families, all belonging to the *Bunyaviricetes* class of viruses. To investigate their TNTase activity, we incubated the L proteins from each virus with their 3′ and 5′ vRNA promoters in presence of each radiolabeled NTP. We found that LACV L is incorporating preferentially ATP in its 3′ promoter and on the 5′/3′ dsRNA promoter instead of the UTP incorporated by HTNV (Figure 7A). As LACV 3′ promoter is able to form a hook and thus a dsRNA (Supplementary Fig. 2), we hypothetize the LACV L also has a TNTase activity that is specific to ATP. When using the 5′ vRNA as substrate, we noticed the multiple incorporation of UTPs. The LACV 5′ vRNA structure predicted (see supplementary figure 2) reveals a possible back-priming event just before multiple adenines near the 3′ end of hook formed by LACV 5′ vRNA (see Table 1). Interestingly, this was not observed with the HNTV polymerase with the same template (see Fig.5 lane 9) suggesting that the capacity of back-priming for an RNA is L protein dependent. CCHFV L protein showed more promiscuity on the specificity of incorporation of NTPs both with the 3′ vRNA, with a preference of incorporation G>U>C>A and an equal incorporation for any NTP on the 3′-5′ vRNA duplex. In this case no NTP incorporation was observed on the 5′ vRNA (Figure 7B).

**Figure 7.**
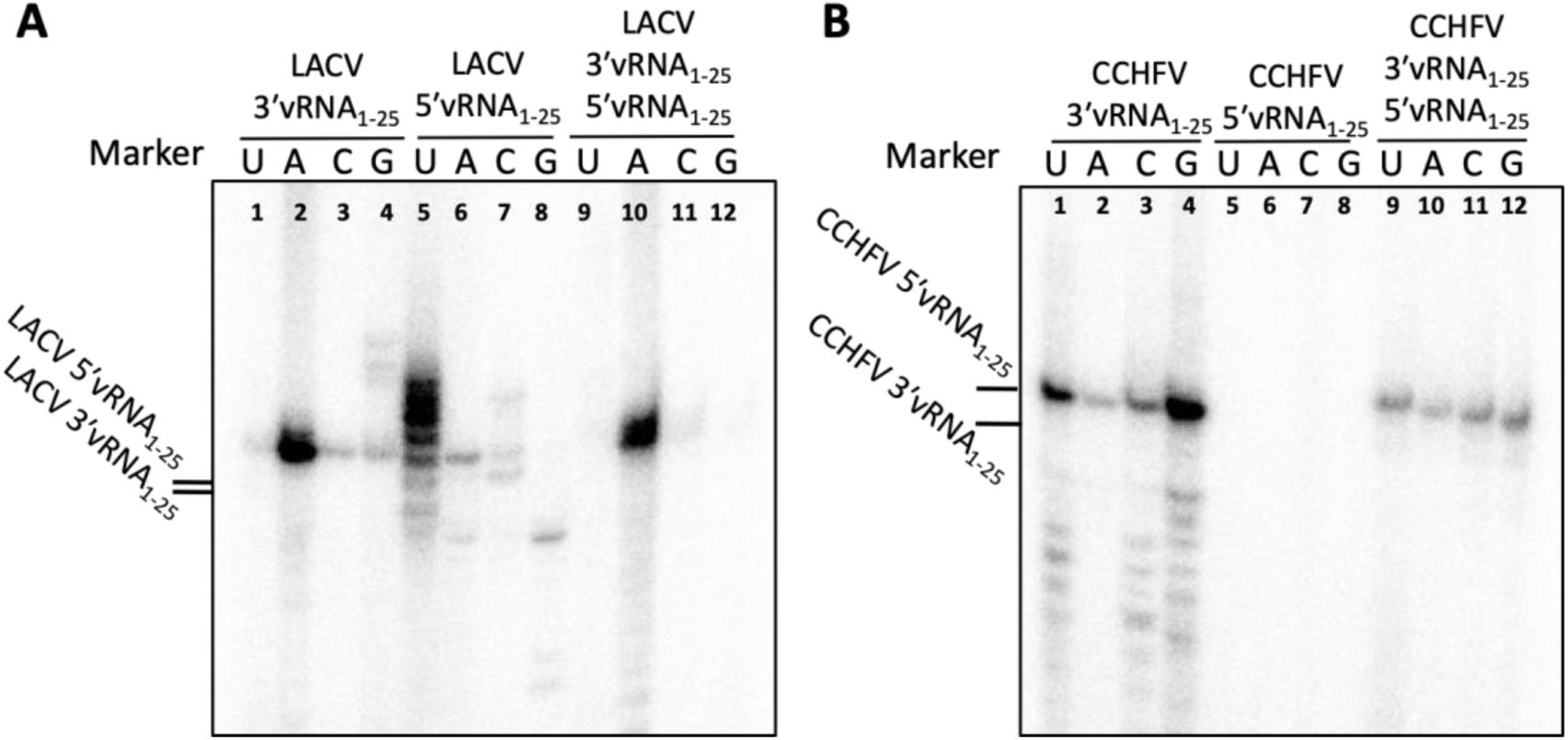
Terminal Nucleotidyl Transferase activity analysis of LACV L_WT_ and CCHFV L_WT_ on their vRNA. (A) TNTase activity of LACV L_WT_ on its vRNA. (B) TNTase activity of CCHFV L_WT_ on its vRNA. Reactions were carried out in the presence of the NTP indicated in each case. Size markers are indicated on the left corresponding to the size of the respective vRNAs labeled with [γ-^32^P]-ATP.

Altogether our observations confirm the existance of a TNTase activity among different families of the *Bunyaviricetes* class of viruses on dsRNAs. However the nucleotide specificity differs between families as well as the ability to incorporate NTPs in their 3′ vRNA templates.

## Discussion

The *in vitro* reconstitution of replication and transcription assays of *Bunyaviricetes* polymerases is paramount for the screening of therapeutic inhibitors for a large set of severe infectious diseases. In this work, we report a new TNTase activity for Hantaan virus, which we also detect on *Peribunyaviridae* and *Nairoviridae* viral families’ L proteins, and therefore points towards being a common feature of sNSVs polymerases. This activity needs to be considered when designing *in vitro* replication assays by including the appropriate controls to avoid misinterpretation of TNTase signal with replication signal as both products migrate at approximately the same positions on chromatography gels. Importantly, TNTase occurs on dsRNA blunt and overhang ends that usually form with *in vitro* incubated WT 3′/5′ Bunyaviral vRNA ends due to their very high complementarity. These conditions favor TNTase activity and do not allow replication activity that necessitates the 10 terminal nucleotides of the 3′vRNA template end to be single-stranded (10). To avoid mis-interpreting TNTase activity with replication activity, and since there is a preference for TNTase nucleotide incorporation for each L protein related to the RNA substrate, we show the need to adequately select the labeled nucleotides used in the replication assays to avoid visualization of TNTase activity or to block the 3′ ends of the template and activator RNA’s by dideoxy terminal nucleotides (Supplementary Fig. 1). Our results show that GTP or ATP, GTP or CTP and ATP appear to be the best nucleotides for replication and transcription assays of Hantavirus, Orthobunyavirus and Nairovirus L proteins respectively, because of the low TNT activity shown with the substrates we use in this study.

TNTase activity is here observed *in vitro* and opens way to, in the future, assess if it exists in the context of the infected cells. During the viral cycle, at the replication termination both 3′ and 5′ are forming a blunt end dsRNA in the L protein active site before splitting, a situation prone for TNT activity. Several hypotheses can be drawn about its potential role(s), that will need to be experimentally confirmed in future studies.

The first proposed potential function is inferred from the TNTase function in positive-stranded RNA viruses. In these viruses, the TNTase activity has been related to the conservation of genome ends. Maintenance of genome ends is essential for the adequate self-recognition of the vRNA and is thus a common issue of living systems. The TNTase activity of Bunyaviruses is intriguing since they have already developed a prime-and-realign mechanism that addresses the genome integrity conservation problem.

One hypothesis explaining the co-existance of both TNT and prime-and-realign mechanism is that since prime-and-realign *de novo* initiation starts in the second or third triplet of the genome end, the addition of nucleotides at the 3′ end may help the polymerase to accommodate the second triplet in the active site if the first triplet is compromised and, in this fashion, contribute to maintain the right genome end in the nascent strand (Fig. 8). This hypothesis needs to be experimentally addressed and, if correct, TNTase activity could thus appear as a versatile and extended feature for RdRps of the viral world.

**Figure 8.**
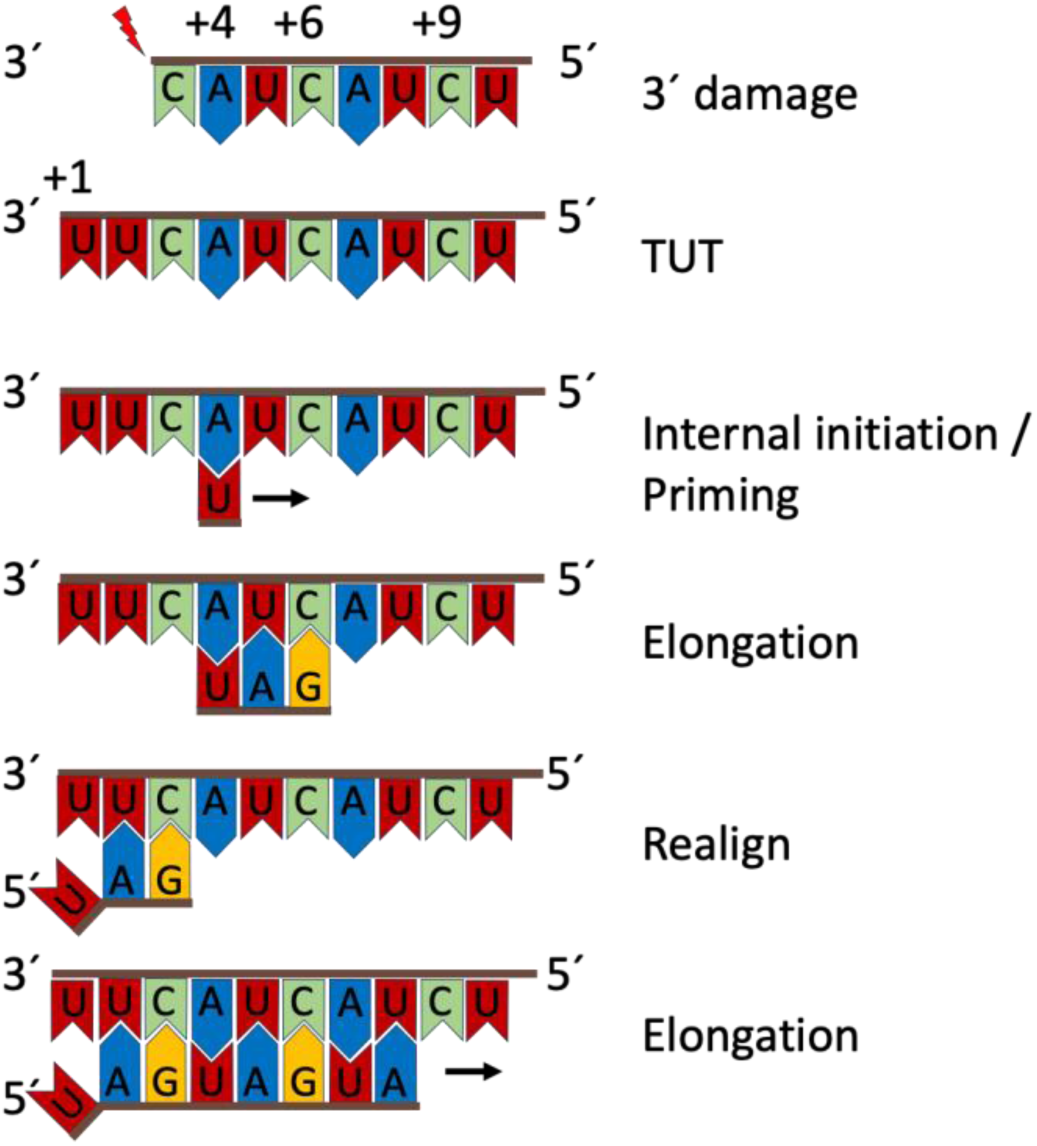
Incorporation of TNT activity in the context of the bunyavirus prime-and-realign initiation mechanism. The addition of nucleotides in the 3′ OH would facilitate the internal initiation in the case of RNA damage, e.g. trimming by cellular endonucleases, and could prevent a blunt end recognizable by RIG-I receptors which would triger interferon response. The newly incorporated TNTs would not be copied to subsequent generations thanks to the prime-and-realign mechanism.

A second hypothesis would be the addition of NTPs at the 3’ end of the genome to prevent the presence of blunt-end double-stranded RNA and avoid interferon response, since blunt ends are specifically recognised by RIG-I cell receptor for triggering interferon response (37), therefore. Our data suggest that addition of NTPs at the 3’ end of the genome could occur at the end of replication, before the separation of progeny and parental strands, which is the moment when the 3′ end is present as dsRNA in the active site. This is consistent with the observation that the presence of free 5′ vRNA activator is not required for the TNTase activity on dsRNA. Addition of non templated NTPs in the 3′ end of the product at this stage would break the blunt end of template/product vRNA pair. This could be important in the event of dsRNA template/product release in order to avoid the activation RIG-I cell receptor and the subsequent triggering of the interferon response.

Beyond the L protein’s role in viral RNA synthesis, the L proteins have access to cellular RNAs in the cell cytoplasm. In this context their TNTase activity may influence RNA metabolism in the cell cytoplasm, spanning the possibilities for host-pathogen interactions and viral control over cell metabolism during infection. RNA metabolism can indeed be regulated by TNTase activity. For instance, studies in *C. elegans* revealed that cellular poly-(A) polymerases (PAPs) are involved in critical processes such as switching from mitosis to meiosis in the germ line by polyadenylation of mRNAs(38). Instead, in fungi *S. pombe* and *A. nidulans* mRNAs are uridylated for subsequent dispatch to degradation (39, 40). PAP GLD-2 in mammals adds a single-AMP which seems to stabilize the miRNA (41). TENT4B polyadenylates human miR-21-5p and enhances its degradation (42, 43). miRNAs metabolism is important to understand infection, they are known to regulate innate immune signaling through IFN production. miRNA expression changes upon activation of innate immune pathways (44) and can directly interfere with viral infection by the targeting of viral RNAs. The crosstalk between infection and miRNAs metabolism has been extensively reported for RNA and DNA viruses (45–49). The intrinsic differences on RNA substrate preference and TNTase nucleotide preference suggest that each viral family could tailor the interactions with cell RNAs differently attending to different infection strategies. Further research is required to determine the effect of the L proteins TNTase activity that we report here on the miRNA cell metabolism.

During the preparation of this article, several studies have seen the light reporting the structure and function of the CCHFV L EN and full length (50–54). The activity of the endonuclease is reported as well as the RNA dependent RNA polymerase activity (53, 54). The EN of CCHVF L appears here as inactive, however experiments in FL-L protein and EN domain isolated show EN activity using different experimental conditions and with no other ENs as reference. This article enriches this evidences by comparison of His (+) and His (-) ENs which shows His (+) ENs much more active than His (-), as previously shown for Arenavirus ENs (20). Replication assays were also reported through different strategies. Replication activity was reported when mutating the RNAs to prevent double stranded formation (50, 54), in other cases activity consistent with TNT activity is reported as RNA synthesis activity (52). These studies thus give new insights on L protein structure and function complementary to the TNTase activity reported in here for bunyaviruses.

## Supplementary Information

**Supplementary Figure 1.**
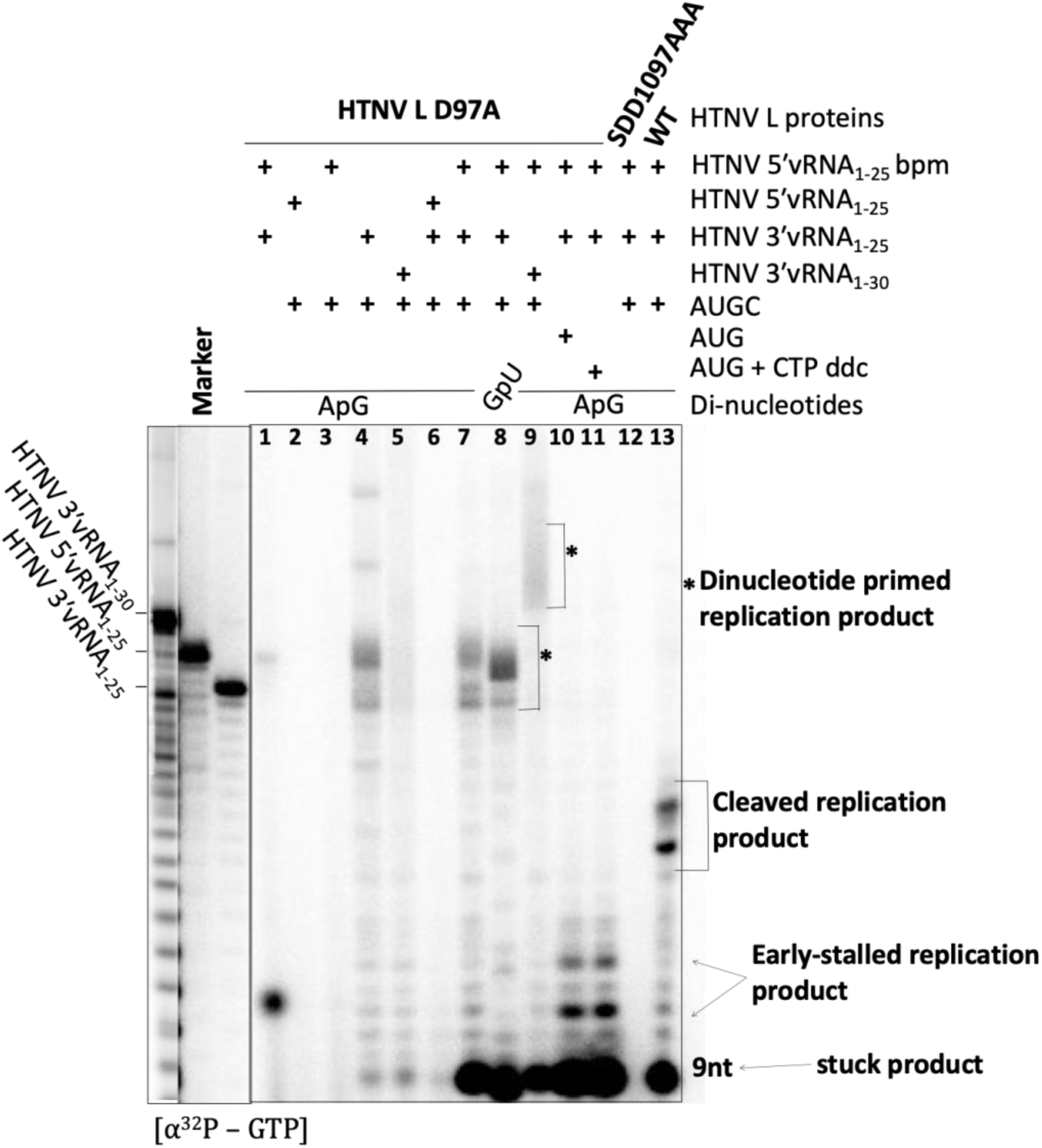
*In vitro* RNA replication activity of HTNV L_WT_, HTNV L_D97A_ and RdRp active site mutant HTNV L_SDD1097AAA_.

*In vitro* replication assay using conditions similar as in (10). This control experiment shows the replication activity of HTNV L. The RNA markers used are 30 and 25 nucleotides-long labelled HTNV vRNA promoters (3′ and 5′). HTNV L needs dinucleotides for *in vitro* replication activity enhancement, the 3′ vRNA template and the 5′ vRNA allosteric regulator. The nucleotides added in the reaction are indicated (AUGC for Adenine, Uridine, Guanine and Citidine) Without the allosteric regulator the enzyme shows some basal activity (lane 4) which is blocked when the 5′ is added to the reaction (lane 6). The reason is that the 5′ and 3′ are almost perfectly complementary, generating a dsRNA that can be uridylated by HTNV L through the TUTase activity. Here we do not see this activity because we are using GTP as labelled nucleotide. To observe the allosteric regulation of the L protein we need to break the complementarity through mutagenesis (5′ base-pair mutation, bpm), in this case we see the synthesis of the RNA of different sizes (lines 7-9) with a characteristic stacked product of about 9 nucleotides. Dinucleotide ApG hybridizes in the first nucleotide and GpU in the second (lanes 7-8). Replication is truncated by the absence of CTP or the presence of ddCTP yielding 10 or 12 nucleotide-long truncated products (lanes 10-11), and abolished by the mutation of the L protein active site (lane 12). The products of replication are cleaved if the endonuclease domain is not inactivated by mutagenesis (D97A) despite the nascent RNA is not capped. This suggests that an in trans endonuclease cleavage is occurring in our experimental conditions, which are far from the context of the ribonucleoprotein during infection (lane 13).

**Supplementary figure 2:**
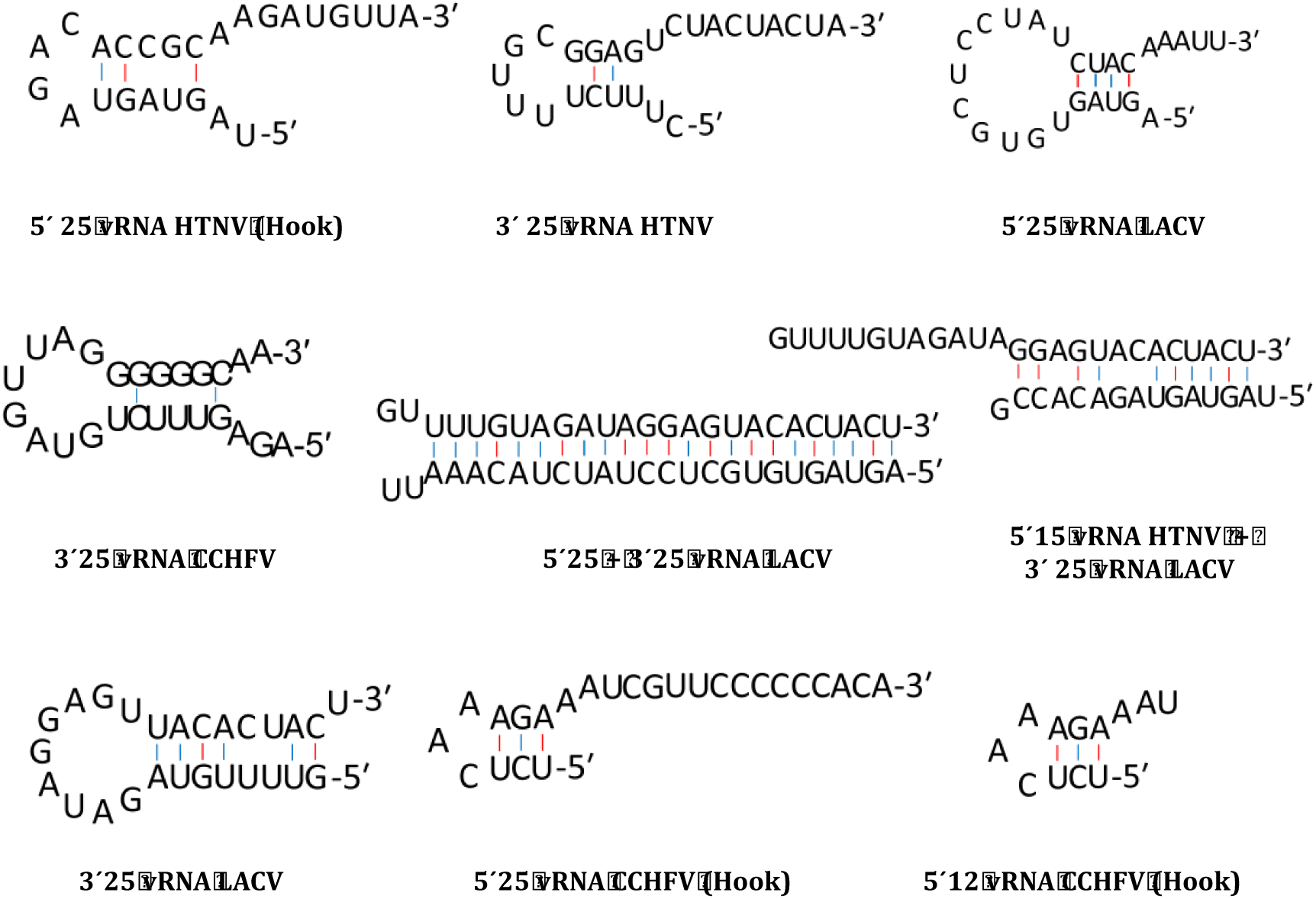
RNA folding. The RNA fold predictions of the RNAs used in this study were carried out using Mfold within UNAfold package web server. Computer-predicted RNA fold was carried out using the default parameters and output RNA secondary structures were consider to create the RNA fold figures.

